# Isoform- and ligand-specific modulation of the adhesion GPCR ADGRL3/Latrophilin3 by a synthetic binder

**DOI:** 10.1101/2022.07.20.500857

**Authors:** Szymon P. Kordon, Przemysław Dutka, Justyna M. Adamska, Sumit J. Bandekar, Katherine Leon, Brock Adams, Jingxian Li, Anthony A. Kossiakoff, Demet Araç

## Abstract

Adhesion G protein-coupled receptors (aGPCRs) are cell-surface proteins with large extracellular regions that bind to multiple ligands to regulate key biological functions including neurodevelopment and organogenesis. Modulating a single function of a specific aGPCR isoform while affecting no other function and no other receptor is not trivial. Here, we engineered an antibody, termed LK30, that binds to the extracellular region of the aGPCR ADGRL3, and specifically acts as an agonist for ADGRL3 but not for its isoform, ADGRL1. The LK30/ADGRL3 complex structure revealed that the LK30 binding site on ADGRL3 overlaps with the binding site for an ADGRL3 ligand – teneurin. In cellular-adhesion assays, LK30 specifically broke the trans-cellular interaction of ADGRL3 with teneurin, but not with another ADGRL3 ligand – FLRT3. Our work provides proof of concept for the modulation of isoform- and ligand-specific aGPCR functions using unique tools, and thus establishes a foundation for the development of fine-tuned aGPCR-targeted therapeutics.

## INTRODUCTION

With 33 members in the human genome, adhesion G protein-coupled receptors (aGPCRs) represent the second-largest subfamily of GPCRs. Genetic studies have identified critical roles for aGPCRs in development, immunity, and neurobiology (1–7) linking them to numerous diseases including neurodevelopmental disorders, deafness, male infertility, schizophrenia, immune disorders, and cancers (2, 8–15). While aGPCRs are crucial surface receptors involved in numerous physiological processes (16), establishing an understanding of their mechanisms of action at the molecular level remains an ongoing challenge, as has been the development of tools to more effectively study them and to modulate their functions.

In addition to their signaling seven transmembrane (7TM) helices that exist in all GPCRs, a hallmark of aGPCRs is their large multidomain extracellular regions (ECRs) (17–19). The multidomain ECRs of aGPCRs can bind to protein ligands that are either on neighboring cells or in the extracellular matrix, effectively regulating receptor function and downstream signaling (11, 20–27). The ECRs of aGPCRs range from ∼200 to ∼5900 amino acids and can be comprised of combinations of approximately 30 different fadtypes of adhesion domains that include the GPCR Autoproteolysis Inducing (GAIN), Epidermal Growth Factor (EGF), lectin (Lec), immunoglobulin (Ig), cadherin, olfactomedin (Olf), pentraxin, laminin and other domains. While the GAIN domain is conserved in aGPCRs, other extracellular domains vary between aGPCRs and enable each receptor to bind specifically to unique ligands and thus mediate different biological functions.

Several studies, including ours, have identified ECR-targeted synthetic proteins that activate or inhibit aGPCRs (22, 28, 29). However, strategies for modulating particular aGPCR-ligand interactions and specific isoforms are not tested and further work for fine-tuning receptor functions is needed. As most aGPCRs mediate multiple functions, targeting only a single function of the receptor, while leaving the other functions unaffected can be challenging. The supposition of our work is that antibody-like molecules that target the ECRs of aGPCRs may result in highly specific functional modulators because aGPCR ECRs are much more diverse than their 7TMs (30). This approach can also regulate specific aGPCR-related activities by targeting the specific aGPCR-ligand interaction that is responsible for the particular activity. In addition, aGPCRs have been shown to have some roles that depend only on their ECR, making them independent of their TM region (31–33). In such cases, ECR-targeted reagents is the only way to modulate these receptor induced activities.

Latrophilins (ADGRLs, LPHNs) constitute a model aGPCR subfamily that play crucial roles in embryogenesis, tissue polarity, and synapse formation; their mutations are associated with numerous cancers and attention deficit-hyperactivity disorder (ADHD) (34–41). There are three ADGRL isoforms (paralogs) in vertebrates (ADGRL1-3). ADGRL1 and ADGRL3 are primarily expressed in the brain, whereas ADGRL2 is ubiquitously expressed (42, 43). The distribution of the ADGRL isoforms in different tissues suggests that each may contribute to a different set of diverse processes. Studies in rats showed that ADGRL3 knockout (KO) results in hyperactivity and increased acoustic reactivity; protein levels of ADGRL1 and ADGRL2 isoforms remained unaltered, showing no compensatory upregulation in ADGRL3 KO rats (44). A recent study in mice showed that the ADGRL2 and ADGRL3 isoforms mediate the formation of distinct synapses on the same hippocampal neuron and cannot compensate for each other suggesting each isoform has distinct functions, which helps explain the specificity of synaptic connections (5, 37, 45). Furthermore, ADGRL isoform-specific remodeling of the actin-associated complexes in HEK293T cells was reported in response to teneurin binding (46). Thus, affinity reagents that target inhibition of isoform-specific interactions are desirable as they will be able to modulate different ADGRL functions.

ADGRL ECRs are comprised of a Lectin (Lec) domain, an Olfactomedin (Olf) domain, a stalk-like region, followed by a Hormone Binding (HormR) domain and a GAIN domain which directly precedes the seven-transmembrane helix region (7TM) (Figure 1A) (19, 20). ADGRLs have numerous endogenous ligands and likely have more unknown ones that will be revealed with further study. Most ligands interact with all three ADGRL isoforms; however, others bind preferably to one of the isoforms (47). Some of the most highly studied ADGRL ligands include teneurins (TENs) (35, 39, 40, 48) and fibronectin leucine-rich repeat transmembrane proteins (FLRTs) (23, 49), although other less studied interactions with neurexins (NRXs) (50) and contactins (51) have also been reported. ADGRL interactions with both TENs and FLRTs have been shown to induce excitatory synapse formation and specification (39, 40). The excitatory synapse formation depends on both TEN and FLRT interactions with ADGRL, although each ligand might be important for different functions. Thus, molecules that can inhibit the interactions between ADGRL isoforms and their ligands are desirable to regulate ADGRL functions more specifically.

**Figure 1.**
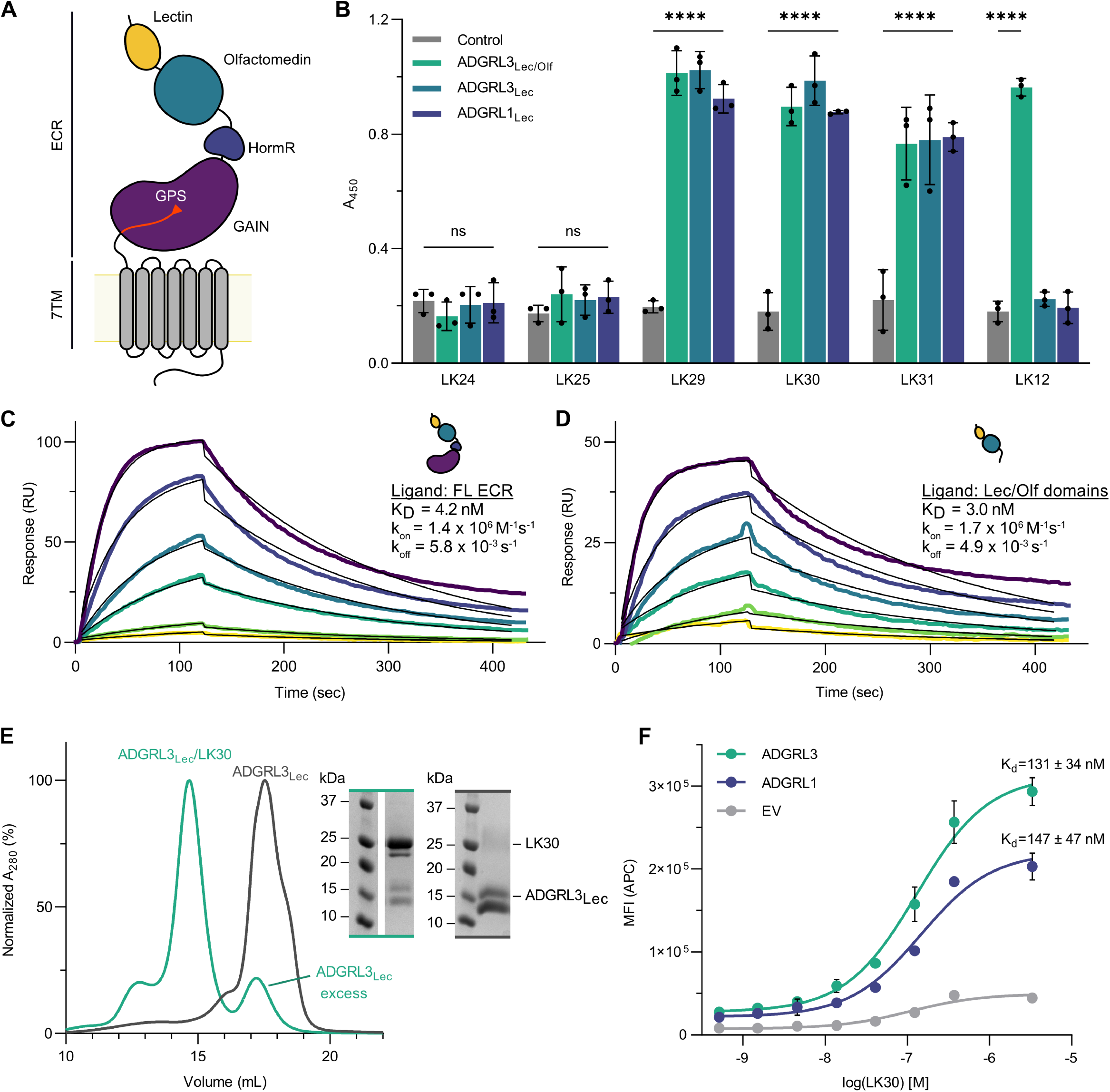
Characterization of the sABs against ADGRL3. **A**. Schematic diagram of full-length human ADGRL3. The Lectin domain is colored yellow, Olfactomedin - cyan, Hormone Binding Region - navy, GAIN - purple and 7TM - gray. The autoproteolysis site within the GAIN domain (GPCR Proteolysis Site (GPS)) and the last β -strand of the GAIN domain is colored red. **B**. Representative single-point protein ELISA of the antibody binders obtained from phage display. Epitope mapping shows that sABs LK27-31 bind to the lectin domain of both ADGRL3 and ADGRL1. Error bars indicate SD (n = 3), ****p < 0.0001 vs. buffer treatment; two-way ANOVA. **C**. and **D**. Surface plasmon resonance measurements of the LK30 binding to purified ADGRL3 ECR (**C**) and Lec/Olf fragment (**D**). Each sAB concentration is shown in a different color trace. Within each plot, the multiconcentration global fit line is shown in black. In order from highest to lowest, the concentrations of analyte used were 25, 12.5, 6.25, 3.125, 1.563, 0.78 1 nM. **E**. SEC profiles and SDS-PAGE analyses show that the LK30 forms a monodisperse complex with the lectin domain of ADGRL3. **F**. Binding activity of LK30 to the receptor was measured using HEK293T cells expressing full-length ADGRL3 (cyan curve) or full-length ADGRL1 (purple curve) by flow cytometry. K_D_ values of LK30 binding were determined as 131 nM and 147 nM, for ADGRL3 and ADGRL1, respectively. Cells transfected with empty vector were used as negative control (gray curve). Error bars indicate SD (n = 3).

In this paper, we used ADGRLs as an exemplary aGPCR system to demonstrate that the ECRs of aGPCRs can be targeted by synthetic binders in a ligand- and isoform-specific manner. Employing phage-display technology, we have generated and characterized a synthetic antibody fragment (sAB) against the ADGRL3 ECR that targets a single domain of ADGRL3. Cell-based signaling assays showed that this sAB acts as an agonist for ADGRL3, but not for ADGRL1, although it binds to both with similar affinities. The crystal structure of the sAB in complex with the Lec domain of ADGRL3 showed that the sAB overlapped with the TEN2 binding epitope on ADGRL3. Herein, we show that specifically breaking ADGRL3’s interaction with one ligand – TEN2 – still allows maintaining the interaction with another – FLRT3. In this work, we have developed valuable tools that will enable further studies of ADGRL function and provide the principles for fine-tuned modulation of aGPCR signaling and downstream biological function.

## RESULTS

### A High-Affinity Domain-Specific Antibody Directed to the Extracellular Region of ADGRL3

In order to generate high-affinity sABs against ADGRL3, biotinylated full-length ECR of ADGRL3 was subjected to phage display selection using a high-diversity synthetic phage library based on a humanized antibody Fab scaffold (52). To increase the specificity and affinity of the sABs, four rounds of selection were performed, with decreasing concentrations of target in each round (Figure S1A). After selection, 96 binders were screened using a single-point phage enzyme-linked immunosorbent assay (ELISA) (Figure S1B). The clones showing high ELISA signal intensity, when compared to control wells were sequenced, identifying a total number of 10 unique binders against ECR of ADGRL3. Selected phagemids were then cloned into sAB protein format, expressed in *E. coli* and purified by protein L affinity chromatography for further characterization.

To determine the epitope of the selected sABs on ECR of the ADGRLs, we performed a single-point protein ELISA, utilizing fragments of either human ADGRL3 or rat ADGRL1 (Figure 1B). Epitope mapping experiments revealed that three out of six ADGRL3 sABs bound to the N-terminal lectin (Lec) domain of the receptor, while one sAB recognized the Olf domain. Interestingly, we also observed that all of the ADGRL3 Lec domain binders can also recognize the Lec domain from ADGRL1.

We determined the binding kinetics of the sABs to their targets by surface plasmon resonance (SPR) (Figure 1C, D, S1C). The binding constants (K_D_) for all of these sABs are in low nanomolar (nM) range with most being characterized by slow dissociation rates (k_off_ < 10^−3^ s^−1^). From the most promising cohort, we focused on the best expressing sAB, LK30, which binds to the ADGRL3 ECR with 4.2 nM affinity for future experiments.

Binding of LK30 to the ECR fragments (Lec and Lec-Olf) of ADGRL3 in solution was confirmed by size exclusion chromatography (SEC). There was a shift in the retention volume of the ADGRL3/LK30 complexes compared with the purified ADGRL3 protein fragments alone and we observed co-elution of both proteins, as analyzed by SDS-PAGE (Figure 1E, S1E). In order to test whether the sABs that were selected against the ECR of ADGRL3 can also bind to full-length ADGRLs expressed on the cell surface, we utilized flow cytometry experiments (Figure 1F, S2A). We expressed either full-length ADGRL3 or ADGRL1 constructs in HEK293T cells and added increasing concentrations of LK30. To detect LK30 binding, we utilized a fluorescently labeled anti-human IgG sAB-fragment specific antibody. For each LK30 concentration, the mean fluorescent intensity (MFI) for 10,000 cells was measured and these values were plotted against the concentration of LK30 to estimate an apparent affinity. We determined that LK30 binds to the full length ADGRL3 and ADGRL1 expressed on the cell surface-expressed receptors with high affinity (131 nM and 147 nM respectively), indicating that its binding epitope is not hindered by the proximity of the membrane or from possible differences in glycosylation pattern in mammalian cells.

### LK30 specifically modulates downstream signaling of ADGRL3, but not ADGRL1

Previous work had shown that the binding of biological ligands to the ECR of other GPCRs can alter receptor signaling ability (22, 28, 29). ADGRL3 signals through Gα_12/13_, Gα_i_ or Gα_q_ proteins, with Gα_12/13_ being activated the most (41, 53). We have previously reported that ADGRL3 and ADGRL1 are active in a serum response element (SRE)-luciferase assay (41). Therefore, we aimed to test the effect of sAB binding on ADGRL3 activity, using an SRE-luciferase assay that measures the receptor signaling. LK30 treatment of ADGRL3 transfected cells resulted in increased signaling of the receptor, as seen on SRE assay results (Figure 2A). The LK30 effect was specific to ADGRL3, as cells transfected with empty vector did not show any significant change in signaling. Treatment with 1 µM of LK30 increased ADGRL3 signaling with an EC_50_ of 48 nM. This was an approximately 2.5-fold increase in signaling compared to the basal activity of the receptor (Figure 2B). Interestingly, we observed no effect of LK30 addition on ADGRL1, even at higher LK30 concentrations (up to 10 µM), despite its ability to bind to ADGRL1 (Figure 2A, B). Thus, these results provide evidence for isoform-specific modulation of ADGRLs by the synthetic ligand LK30.

**Figure 2.**
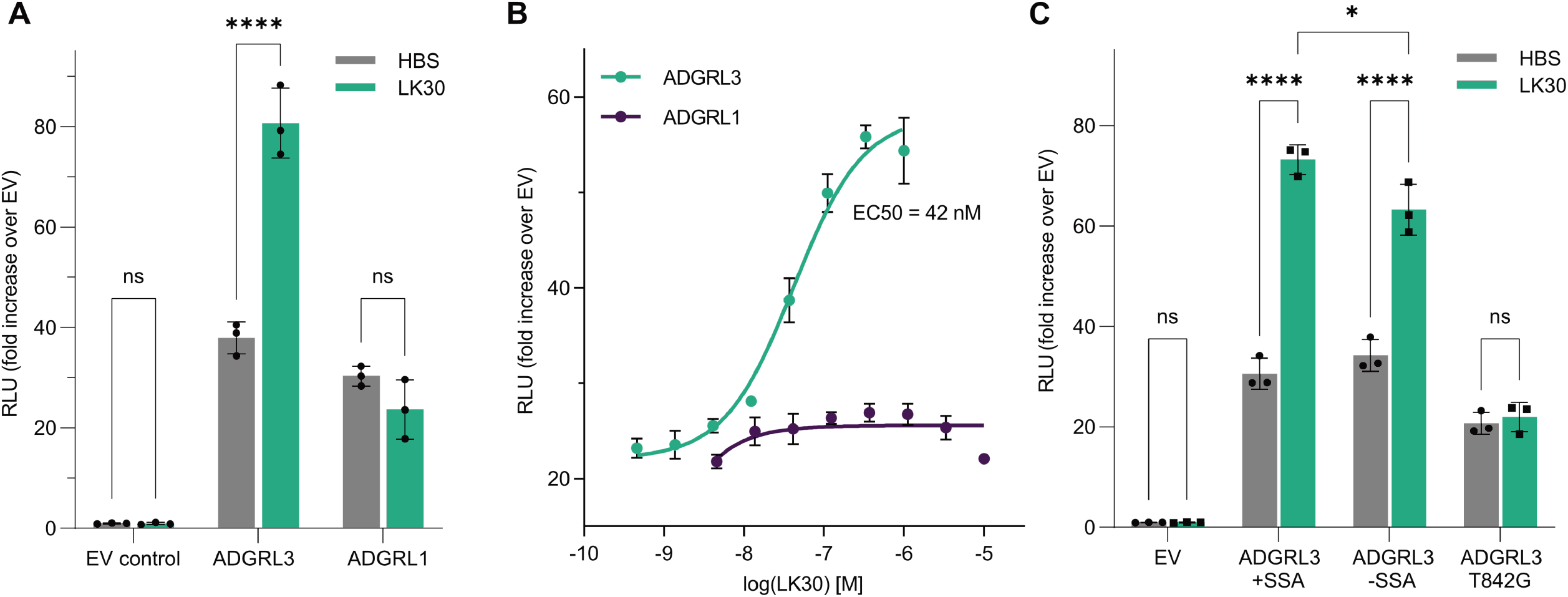
LK30 is an ADGRL3-specific activator. **A**. SRE-luciferase assay for signaling of ADGRL3 and ADGRL1 in the absence or presence of 1 µM purified LK30 presented as fold increa se over empty vector (EV). RLU, relative luminescence units. Error bars indicate SD (n = 3), ****p < 0.0001 vs. buffer treatment; two-way ANOVA. **B**. Titration of LK30 on SRE-luciferase activity of ADGRL3 (cyan) and ADGRL1 (purple). The EC _50_ value of LK30 on ADGRL3 was determined to be 42 nM. Error bars indicate SD (n = 3) **C**. SRE-luciferase assay for the splice isoforms of ADGRL3 (+SSA and -SSA) and ADGRL3 autoproteolysis-null mutant (T842G, +SSA) RLU, relative luminescence units. Error bars indicated SD (n = 3), ****p < 0.0001 vs. buffer treatment; two-way ANOVA.

Previously, we have reported basal activity of ADGRLs in the GloSensor assay (Promega), which reports increase or decrease of intracellular cAMP levels in mammalian cells (26, 41). We have shown that cAMP levels in cells expressing ADGRL3 were significantly lower compared to control. Therefore, we have tested the ADGRL3-activating LK30 in the cAMP assay. In contrast to the SRE-luciferase assay, LK30 showed no significant effect on ADGRL3-dependent cAMP levels (Figure S3).

### Mechanism of ADGRL3 activation by LK30 is autoproteolysis dependent

aGPCRs are cleaved in the conserved GAIN domain by an autoproteolytic mechanism (20). Upon cleavage, the two fragments remain tightly associated (20). Recent studies proposed two complementary models for modulation of aGPCR activity by ligands. In the Stachel-dependent model, ECR dissociation exposes the tethered agonist peptide (the Stachel peptide), allowing its direct interaction with 7TM domain and stimulating receptor activity (54–56). GAIN domain autoproteolysis plays a crucial role in this model, allowing for the possible ECR dissociation. In the Stachel-independent model, ligand interaction with ECR induces conformational changes, allowing for direct and transient interaction between the ECR and 7TM domain (21, 22, 24). Contrary to Stachel-dependent model, regulation occurring through this mechanism of aGPCR activation is independent of the receptor autoproteolysis.

We introduced a single-point mutation, T842G within the GAIN domain of ADGRL3, in order to establish whether LK30 activates ADGRL3 in an autoproteolysis-dependent or -independent manner. We had previously shown that T842G mutation abolishes autoproteolysis within the GAIN domain without disrupting folding or cell-surface trafficking of the mutant receptor (20). Using the SRE-luciferase assay, we first found that the basal activity of the autoproteolysis mutant is not affected when compared to the wild type receptor (Figure 2C). LK30 treatment increased basal activity of ADGRL3 in HEK293T cells transfected with wild type ADGRL3; however, the effect of LK30 was nearly abolished when the cells were transfected with the ADGRL3 T842G mutant (Figure 2C). This result suggests that the agonistic effect of LK30 on the receptor signaling is dependent on the ADGRL3 autoproteolysis.

The ECR of ADGRL3 has a short alternatively-spliced region between Lec and Olf domains (a five amino acid splice insert: KVEQK) that was reported to decrease the affinity of ADGRLs to TENs when present (48). Previous work had shown the importance of alternative splicing in regulating protein-protein interactions and functions (39)(27). In order to test the possible regulation of the LK30 effect on ADGRL3 signaling by alternative-splicing, we designed a ADGRL3 construct removing the five amino acids insert between Lec and Olf domains (ADGRL3 -SSA) and compared it to the construct that we have used throughout this study, ADGRL3 +SSA. LK30 treatment increased signaling of ADGRL3 -SSA with similar fold increase to the wild type ADGRL3 +SSA isoform, suggesting that the LK30 activation of ADGRL3 is not dependent on the splice isoform of the receptor (Figure 2C).

### Crystal structure of ADGRL3/LK30 complex

To elucidate the molecular basis of the interaction between LK30 and ADGRL3 ECR, we determined the crystal structure of the Lec domain of ADGRL3 (ADGRL3_Lec_) in complex with LK30 at 2.65 Å resolution (Figure 3A, S4A and Table 1). The crystal contacts in the structure are mediated predominantly by the heavy chain (HC) and light chain (LC) of the LK30, as well as by the Lec domains (Figure S4B). LK30 binds to Lec through CDRs in both HC (H1, H2, H3) and LC (L3), resulting in the total interface area of 608 Å^2^ (HC – 558 Å^2^; LC – 50 Å^2^) in the protein complex. This interface is mediated by aromatic, and hydrophobic residues of LK30 CDRs and involves extensive hydrogen bonding and van der Waals interactions with residues in β1-β2 and β3-H1 loops at the tip of the Lec domain (Figure 3B). Notably, hydrogen bonds between E37_Lec_ and Y34_LK30_, Y39_Lec_ and Q106_LK30_, D67_Lec_ and Y102_LK30_, K115_Lec_ and Y57_LK30_ and Q72_Lec_ and S55/S58_LK30_ appear to stabilize the interaction and shape the total buried surface area.

**Figure 3.**
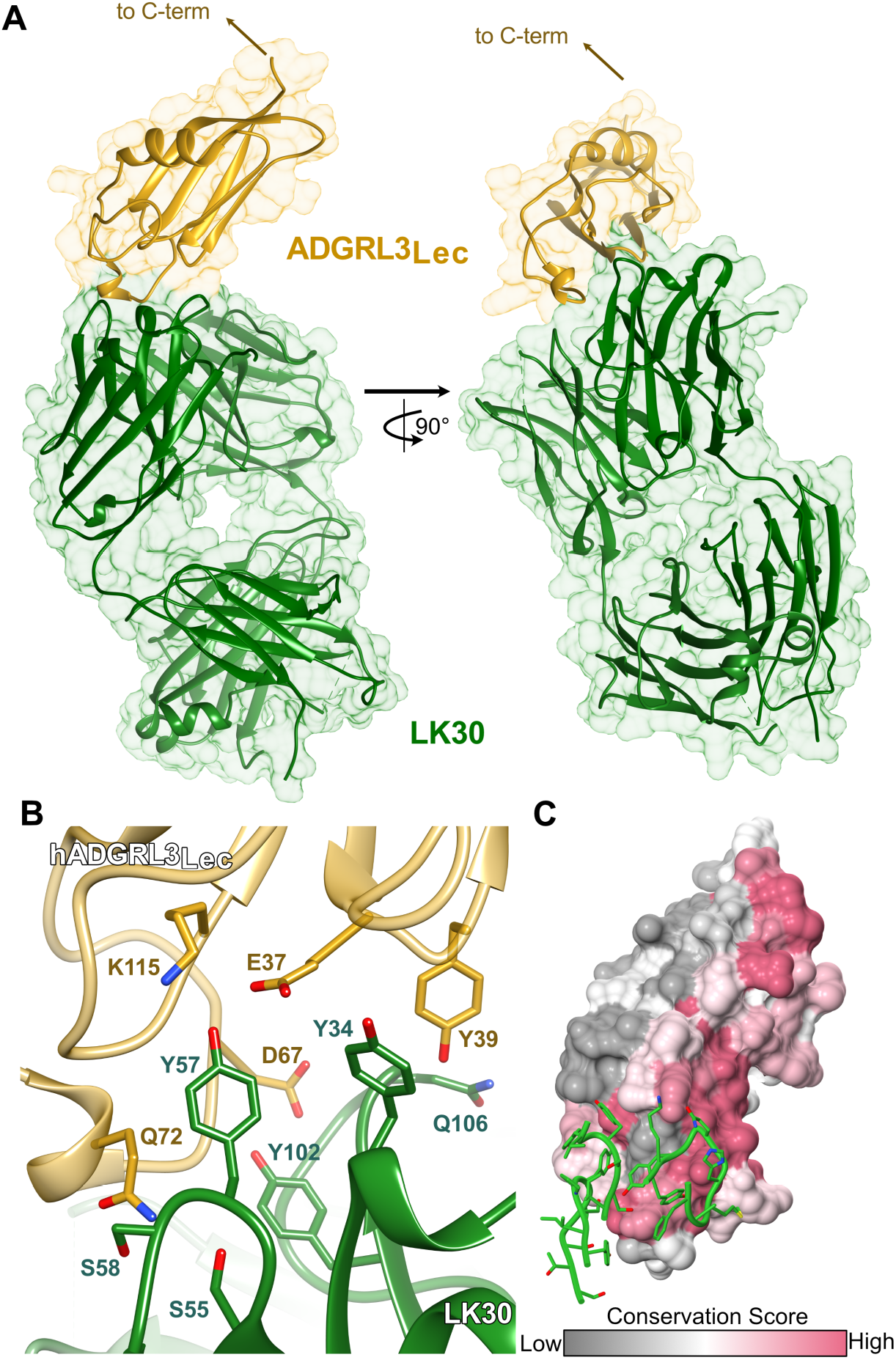
Crystal structure of the LK30/ADGRL3 complex at 2.65 Å resolution. **A**. The crystal structure of the ADGRL3 lectin domain in complex with LK30. ADGRL3Lec is colored yellow while the HC and LC of LK30 are colored green. **B**. Close-up view of the ADGRL3Lec/LK30 interface. Residues at the binding interface are shown as sticks. ADGRL3Lec is colored yellow while the LK30 is colored green. **C**. Surface conservation analysis (gray, variable; red, conserved) of the ADGRL3 Lec domain. The CDRs of the LK30 HC interacting with the Lec domain are shown in green.

**Table 1.** Data collection and refinement statistics.

| **ADGRL3<sub>Lec</sub>/LK30** |  |
| --- | --- |
| **Data Collection Statistics** |  |
| **Wavelength (Å)** | 1.033 |
| **Resolution range (Å)** | 47.51 - 2.33 (2.40 - 2.33) |
| **Space group** | P 21 21 21 |
| **Unit cell dimensions (Å, °)** | 81.80 97.85 163.06 90 90 90 |
| **Total reflections** | 108040 (8833) |
| **Unique reflections** | 56710 (4563) |
| **Multiplicity** | 1.9 (1.9) |
| **Completeness (%)** | 100 (99.9) |
| **Mean I/sigma(I)** | 9.3 (0.1) |
| **Wilson B-factor (1/Å<sup>2</sup>)** | 80.04 |
| **R-merge** | 0.038 (7.31) |
| **R-meas** | 0.054 (10.34) |
| **R-pim** | 0.038 (7.31) |
| **CC1/2** | 0.99 (0.079) |
| **CC\*** | 1 (0.38) |
| **Refinement Statistics** |  |
| **Resolution range (Å)** | 47.51 - 2.65 (2.74 - 2.65) |
| **Reflections used in refinement** | 38700 (3768) |
| **Reflections used for R-free** | 3521 (352) |
| **R-work** | 0.24 (0.38) |
| **R-free** | 0.27 (0.40) |
| **CC (work)** | 0.94 (0.63) |
| **CC (free)** | 0.92 (0.52) |
| **Number of non-hydrogen atoms** | 8099 |
| **Macromolecules** | 8058 |
| **Ligands** | 34 |
| **Solvent** | 7 |
| **Protein residues** | 1059 |
| **RMS (bonds) (Å)** | 0.012 |
| **RMS (angles) (°)** | 1.56 |
| **Ramachandran favored (%)** | 95.39 |
| **Ramachandran allowed (%)** | 4.51 |
| **Ramachandran outliers (%)** | 0.10 |
| **Rotamer outliers (%)** | 1.86 |
| **Clashscore** | 7.74 |
| **MolProbity Score** | 1.95 |
| **Average B-factor (Å<sup>2</sup>)** | 104.6 |
| **Macromolecules** | 104.6 |
| **Ligands** | 114.79 |
| **Solvent** | 72.43 |
| **Number of TLS groups** | 29 |

### LK30 specifically breaks the interaction of ADGRL3 with TEN2, but not FLRT3

To visualize conserved and variable regions of the ADGRL3 Lec domain, we analyzed a heat map based on sequence conservation displayed (colored from most to least conserved) on the ADGRL3_Lec_/LK30 complex structure (Figure 3C). We then mapped CDRs of the LK30 sAB (green sticks) on the ADGRL3_Lec_ surface. Sequence conservation analysis revealed that LK30 binds to a highly conserved region on the N-terminal part of the Lec domain. As the ECR of ADGRLs has been previously shown to facilitate the interaction with its endogenous ligands – TENs and FLRTs (Figure 4A), we hypothesized that LK30 binding might prevent those interactions. The structures of ADGRL bound to both TEN and FLRT have previously been reported (23, 39, 40, 57). We superimposed the ADGRL3_Lec_/LK30 complex onto the structures of the ADGRL3/TEN2 and ADGRL3/FLRT3 complexes (Figure 4B) showing the TEN binding site on the Lec domain of ADGRL overlaps almost the entire sAB epitope. A more detailed analysis of the binding interfaces revealed that the LK30 binding site overlaps with the bottom part of the ADGRL/TEN complex interface (Figure 4C). Notably, LK30 binding hinders one of the previously reported salt bridges – between D67 of ADGRL3 and K1712 of TEN2 from forming. Residue S38 of ADGRL3, which has been shown to interact with a conserved N-linked glycosylation on TEN2-N1681, is located in the buried hydrophobic pocket in the LK30 complex structure. Additionally, the LK30 binding blocks the interaction between the Lec domain and residues D1737 and H1738 of TEN2, that has been shown to be crucial for the ADGRL3/TEN2 interaction (39, 40). On the other hand, analysis of the FLRT binding site on the Olf domain of ADGRL3 suggests that LK30 interaction should not affect FLRT binding to ADGRL3. The remaining domains of ADGRL3 face away from the sAB, suggesting that LK30 would only interfere with TEN2 binding, but not with FLRT binding (Figure 4B).

**Figure 4.**
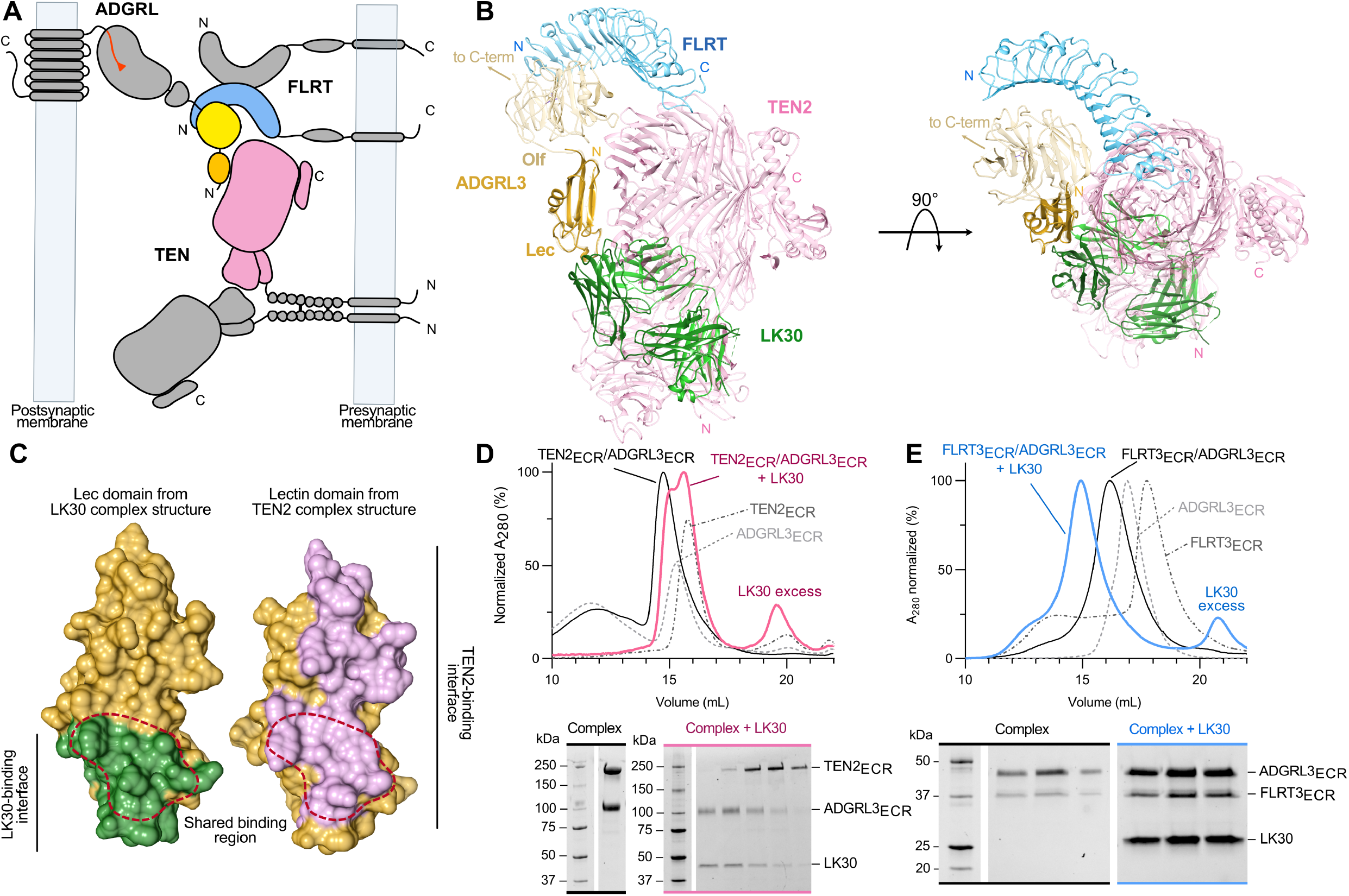
LK30 blocks the interaction of ADGRL3 with TEN2 but not with FLRT3. **A**. Schematic diagram of the interaction network between TEN, ADGRL, and FLRT at the synapse. TEN and FLRT are localized on the presynaptic cell membrane, while ADGRL is localized on the postsynaptic membrane. **B**. Superimposition of the ADGRL3Lec/LK30 complex with the ADGRL1/TEN2 complex structure (PDB: 6SKA) (40) and trimeric complex of ADGRL3 and FLRT2 (PDB: 5FTU) (57). The ADGRL domains of all structures are superimposed. ADGRL3Lec and LK30 are colored yellow and green, respectively, TEN2 molecule is colored pink, FLRT is colored blue. The lectin and olfactomedin domains bound to TEN2 and FLRT2 are colored tan. **C**. Detailed analysis of ADGRL3 lectin domain regions interacting with either LK30 (left, in cyan) or TEN2 (right, in purple). Binding area shared by both ligands on lectin domain is indicated by a red dashed -line. **D**. SEC profiles showing the disruption of the ADGRL3/TEN2 complex (black curve) by the LK30. Addition of the LK30 to the binary ADGRL3 /TEN2 complex leads to TEN2 dissociation and formation of ADGRL3/LK30 complex, as observed on SEC profile (pink curve) and SDS -PAGE analysis. Each chromatogram color corresponds to the accompanying SDS-PAGE gel label. SEC profiles of individual proteins are shown as dashed curves. **E**. SEC profiles showing the formation of the trimeric FLRT3/ADGRL3/LK30 complex (blue curve). Addition of the LK30 to the binary ADG RL3/FLRT3 (black curve) complex causes a shift in the retention volume on SEC profile. SEC profiles of individual proteins are shown as dashed curves. Each chromatogram color matches the accompanying SDS-PAGE gel.

To test whether LK30 can prevent ADGRL/TEN complex formation, we performed SEC experiments with the ECRs of ADGRL3/TEN2. The addition of LK30, resulted in the dissociation of the ADGRL3/TEN2 complex, as observed on SEC chromatogram and SDS-PAGE analysis (Figure 4D). The single peak of ADGRL3/TEN2 complex reforms into the complex of ADGRL3/LK30 (higher order species on SEC curve and first fractions of corresponding gel) and free TEN2 ECR (secondary peak on SEC curve and last three fractions of the gel). We performed a similar set of experiments to test the ADGRL3 binding to FLRT3 (Figure 4E). As expected from the structural analysis, we did not observe ADGRL3/FLRT3 complex dissociation. Instead, after addition of LK30 we observed a further shift in the retention volume, corresponding to the LK30/ADGRL3/FLRT3 trimeric complex formation as confirmed by SDS-PAGE analysis.

### sABs break specific intercellular contacts formed by ADGRL3-TEN2 and ADGRL3-FLRT3 interactions

Cell-adhesion proteins can interact with each other either in *cis*- or *trans*-. When expressed on the same cell surface, cell-adhesion proteins might be involved in *cis*-interactions. Alternatively, when cell-adhesion proteins are expressed on the neighboring cells, they might be involved in *trans*-interactions that mediate cell-cell adhesion and intercellular contacts. Previous studies have shown that ADGRLs interact with TEN2 and with FLRT3 in a trans-cellular manner and promote cell adhesion (23, 48). To examine the effect of LK30 on the interaction of ADGRL3 with its ligands, we performed cell aggregation assays with HEK293 cells in which each full-length protein is expressed on different cell populations and the cells are then mixed to monitor cell adhesion observed as cell-cell aggregation. As expected, when mixing cells expressing ADGRL3 with either TEN2- or FLRT3-expressing cells, we detected formation of significant cell aggregates when compared to control samples (Figure 5A, B and F, G). Addition of LK30 to the mixtures significantly abolished ADGRL3/TEN2-mediated cell adhesion, validating our SEC results in the context of full-length receptors (Figure 5C). LK30 binding had no effect of ADGRL3/FLRT3 interaction and cell aggregation suggesting that LK30 acts specifically on the ADGRL3-TEN2 interaction (Figure 5H). Furthermore, we performed cell aggregation assays with another sAB called LK12, that we characterized as a ADGRL3 Olf domain binder (Figure 1B and S5). As FLRT3 interacts with the Olf domain of ADGRL3, our expectation was that LK12 might specifically affect ADGRL3-FLRT3 interaction. In contrast to LK30, LK12 abolished the ADGRL3 interaction with FLRT3, inhibiting ADGRL3/FLRT3-mediated cell adhesion, but did not affect ADGRL3 binding to TEN2 (Figure 5D, I).

**Figure 5.**
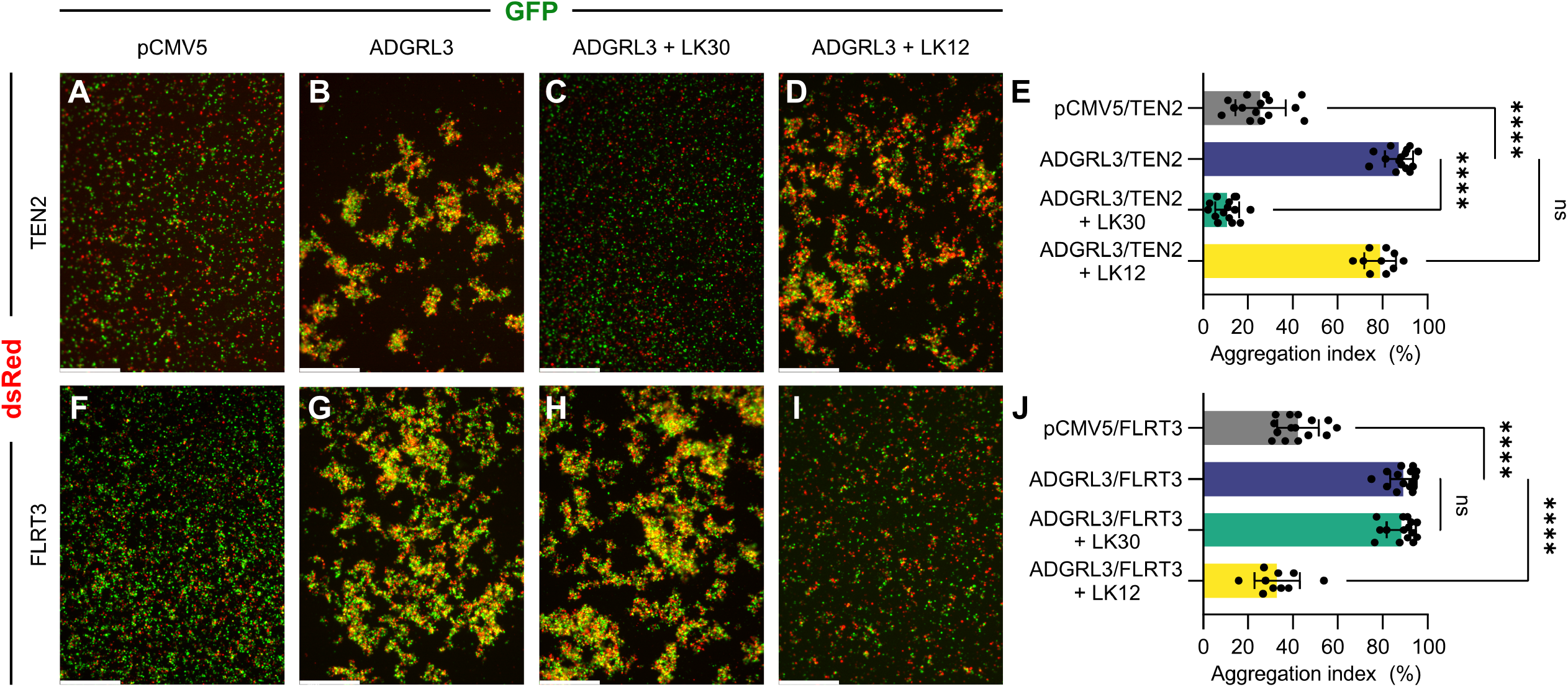
sABs specifically inhibit cell-cell adhesions mediated by ADGRL3-ligand interactions. **A-D**. and **F-I**. Cell-aggregation experiments show full-length ADGRL3 expressed on HEK293T cells interact with full-length TEN2 expressed on another population of HEK293T cells in a trans-cellular manner (**A, B**) Similarly, ADGRL3 interacts with full-length FLRT3 expressed on HEK293T cells in a trans-cellular manner as well (**F, G**). Addition of sAB LK30 breaks the ADGRL3/TEN2 interaction and abolishes cell adhesion (**C**.), but does not interfere with ADGRL3/FLRT3 interaction (**H**.). In contrast, sAB LK12 breaks ADGRL3 interaction with FLRT3 (**I**.), but not ADGRL3/TEN2-mediated cell adhesion (**D**.). HEK293 cells were co-transfected with ADGRL3 or TEN2/FLRT3 and either GFP or dsRed as indicated. Scale bars: 500 μm. **E**. and **J**. Quantification of aggregation index are presented as mean, error bars indicate SD (n = 15 or n = 10 for LK12 experiments), ****p < 0.0001; one-way ANOVA.

Taken together, these data show that we have developed highly specific sAB binders that can specifically block the ADGRL3 interaction with only one of its ligands, while preserving the interaction with the other.

## DISCUSSION

aGPCRs are large chimeric molecules with transmembrane regions that are structurally homologous to the seven-pass GPCRs, and with ECRs that are homologous to cell-adhesion molecules (such as cadherins), receptor tyrosine kinases (such as EGF receptor) and others. Among the 33 human members of aGPCRs, 32 of them have ECRs and comprise numerous extracellular adhesion domains (ranging from one domain in ADGRG1/GPR56 to 40 domains in ADGRV1/GPR98) in addition to their conserved GAIN domain.

In this work, we employed the ADGRL subfamily of aGPCRs as a model system to demonstrate that the ECR of aGPCRs can be specifically targeted by antibodies. Utilizing the phage display selection, we generated a number of ADGRL-specific synthetic antibodies. From this cohort, we focused on characterizing LK30 and showed it to be an activator of ADGRL3-dependent SRE signaling. We determined that LK30 binding to the N-terminal Lec domain of the ADGRL3, distal from the 7TM region, increases the basal signaling of the receptor. To further investigate LK30-dependent stimulation, we tested the effect of autoproteolysis within the GAIN domain on the receptor modulation and found that the cleavage at the GPS site is required for the ADGRL3 activation by LK30. These results, along with our previously published work on Stachel-independent modulation of ADGRG1 signaling (22), present two different mechanisms of ECR-targeted aGPCRs activation.

When targeting receptors, one has to consider different isoforms (paralogs which have been generated as a result of gene duplication) and splice variants of the gene. Each isoform may have evolved to facilitate different functions (58–60) and specific targeting of only one of them can be crucial. Similarly, the importance of alternative splicing in regulating protein-protein interactions and differentiating protein function has been widely studied (1, 27, 39). In this regard, the LK30 Fab shows that antibodies can act in an isoform-specific manner. The SRE-signaling assays revealed that LK30 acts as an agonist only for ADGRL3, but, does not activate ADGRL1, although it can bind to both isoforms with similar affinity (Figure 6A). This further provides an example where antibody-mediated modulation of receptor signaling can be unique for different isoforms. Though the mechanism of the isoform-specificity remains unknown, we speculate that the explicit change of the ECR conformation necessary to allosterically activate the receptor cannot be induced by LK30 for ADGRL1, due to differences in the ECR sequence. In contrast to isoform-specificity, LK30 activation does not depend on ADGRL3 splice site insertion. As for the case of aGPCRs, the specific functions of aGPCR isoforms are still under investigation. However, there is growing evidence that the different ADGRL isoforms are critical for synapse formation at different sublocations within the same neuron (37, 45). LK30 or similarly specific antibodies have the potential to specifically modulate certain types of synapse formation in a single neuron.

**Figure 6.**
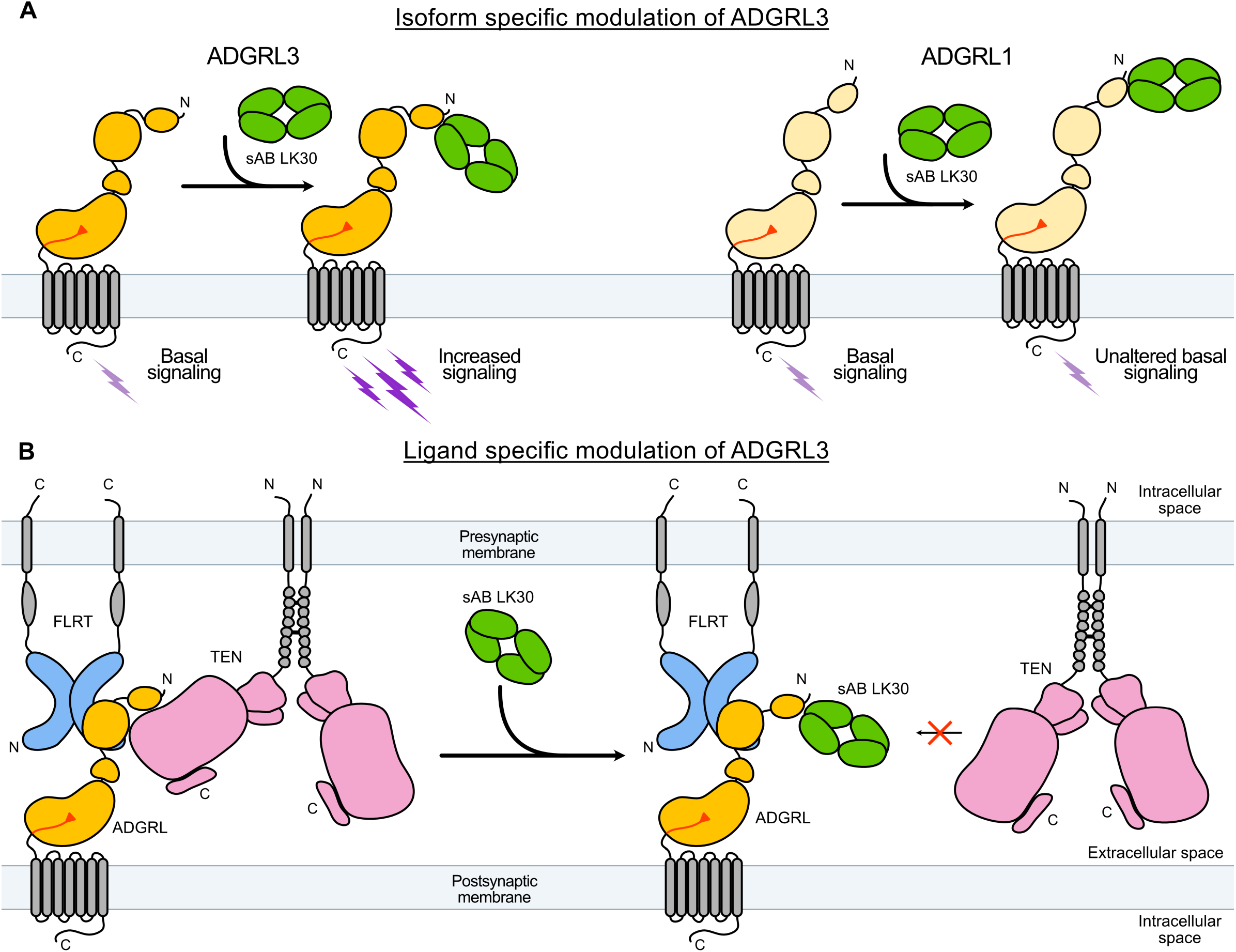
LK30 modulates ADGRL3 in an isoform- and ligand-dependent manner. **A**.Binding of LK30 to ADGRLs modulates the receptor activity in an isoform-specific manner. ADGRL3 (dark yellow) basal signaling increases upon binding of LK30 to the Lec domain of the receptor. The interaction of LK30 with ADGRL1 (light yellow) does not change the signaling activity in the SRE assay. **B**.LK30 breaks the interaction of ADGRL3 with TEN2, whereas it has no effect on the interaction of ADGRL3 with FLRT3. Synthetic antibodies can be used to specifically target and block interactions of the receptor with its endogenous ligand.

The crystal structure of the LK30 in complex with Lec domain of ADGRL3 revealed the LK30 binding site on ADGRL3 surface. Comparison to other available ADGRL complex structures revealed that the LK30 epitope partially overlaps with the TEN binding site. We showed that the LK30 can break the interaction of ADGRL with TEN while maintaining its interaction with FLRT (Figure 6B). Additionally, utilizing another sAB – LK12 that inhibit ADGRL3/FLRT3 complex formation, we demonstrate that synthetic antibodies can be used to specifically target and block interactions of the receptor with only one of its endogenous ligands.

Although most aGPCRs are still orphan receptors with no known ligands, it is reasonable to assume that other aGPCRs also have many ligands that bind to the ECR and modulate receptor function. For instance, the ADGRG1 has three known binding partners, each attributed to its three different functions: the interaction of ADGRG1 with collagen 3 mediates brain cortex development, while tissue transglutaminase 2 mediates central nervous system myelination and phosphatidylserine mediates microglia activation (61, 62). Drugging aGPCRs will thus, require a sophisticated approach rather than simply turning on or off their downstream signaling by targeting the TM. Indeed, it has been reported that the small molecule α-DOG that binds to the 7TM of an ADGRG1 and acts as an agonist, also affects another aGPCR ADGRG5/GPR114 (63). Because of the high variation and diversity of the aGPCR ECRs, the development of ECR-targeted ligands is more likely to result in highly specific reagents for aGPCRs. An additional advantage of targeting the ECR is gaining the ability to disrupt the interactions between aGPCRs and their ligands in a specific manner. Thus, precise targeting of aGPCRs by synthetic molecules in a ligand- and isoform-specific manner may be used as a foundation for drug design to treat aGPCR-mediated diseases.

## METHODS

### Protein expression and purification

The DNA constructs for protein expression and purification were published previously (23, 26, 39). All proteins were expressed using the baculovirus method. *Spodoptera frugiperda* Sf9 insect cells were co-transfected with the constructed plasmids and linearized baculovirus DNA (Expression Systems, 91-002) using Cellfectin II (Thermo Fisher, 10362100). Baculovirus was amplified in Sf9 cells in SF-900 III medium supplemented with 10% FBS (Sigma–Aldrich, F0926).

ECR constructs of ADGRL3, TEN2 and FLRT3 were expressed in High Five cells (Thermo Fisher, B85502). Cell cultures grown in Insect-Xpress medium (Lonza, 12-730Q) were infected with high-titer baculovirus at a density of 2.0 × 10^6^ cells ml^−1^ and incubated for 72 h at 27 °C. The cells were pelleted by centrifugation and the conditioned medium containing the secreted glycosylated proteins were collected. Final concentrations of 50 mM Tris pH 8, 5 mM CaCl_2_, and 1 mM NiCl_2_ were added to the media, the mixture was then stirred for 30 min and centrifuged at 8000 *g* for 30 min to remove the precipitate. The supernatant was incubated with Ni-NTA resin (Qiagen, 30250) for 3 h. The resin was collected with a glass Buchner funnel and washed with 10 mM HEPES pH 7.2, 150 mM NaCl and 20 mM imidazole. Avi-tagged proteins were then biotinylated on-column, incubating the resin-bound proteins with 50 mM bicine pH 8.3, 10 mM MgOAc, 100 mM NaCl, 10 mM ATP, 0.5 mM biotin and 5mM BirA at 27°C with gentle mixing. The protein sample was later eluted with 10 mM HEPES pH 7.2, 150 mM NaCl and 200 mM imidazole. The elution fractions were concentrated and loaded onto a Superdex S200 10/300 GL column (GE Healthcare) equilibrated with 10 mM HEPES pH 7.2 and 150 mM NaCl. Purified fractions of the complex were collected for further experiments.

### Phage Display Selection

Selection for ECR fragments of ADGRL3 was performed according to previously published protocols (52, 64). 200 nM of target was immobilized on streptavidin magnetic beads for the first round of selection. Next, the beads were washed three times to remove unbound protein and 5 mM D-biotin was added to block unoccupied streptavidin on the beads to prevent nonspecific binding of the phage. Afterwards, the beads were incubated for 30 min at RT with the phage library E (65), containing 10^12^-10^13^ virions ml^-1^ with gentle shaking. This was followed by washing of beads containing bound phages, which were later used to infect log phase *E. coli* XL1-Blue cells. Infected cells were grown overnight in 2YT media with 50 µg/mL ampicillin and 10^9^ p.f.u. ml^-1^ of M13 KO7 helper phage in order to amplify phages. Three additional rounds of selection were performed with decreasing target concentration in each round (100 nM, 50 nM, 10 nM) using the amplified pool of virions of the prior round used as the input. Rounds 2 to 4 were performed using semi-automated platform using the Kingfisher instrument. In those rounds phages were eluted using 0.1 M glycine pH 2.7. This technique often risks the enrichment of nonspecific and streptavidin binders. In order to eliminate them, the precipitated phage pool from rounds 2 to 4 were negatively selected against 100 mL of SA beads. The ‘‘precleared’’ phage pool was then used as an input for the selection.

### Enzyme-Linked Immunoabsorbent Assays (ELISA)

ELISA experiments were carried out using a 96-well flat-bottom plate coated with 50 µL of 2 mg ml^-1^ neutravidin in Na_2_CO_3_ pH 9.6 and subsequently blocked with 0.5% Bovine Serum Albumin in 1×PBS. Binding screens of all of the selected sABs in phage format was performed using a single point phage ELISA. 400 µL of 2YT media with 100 µg/ml ampicillin and M13 KO7 helper phage were inoculated with single *E. coli* XL1-Blue colonies harboring phagemids, and cultures were grown at 37 °C for 18h in a 96-deep-well block plate. The cells were pelleted by centrifugation and sAB phage-containing supernatants were diluted 20× in ELISA buffer. Diluted phages were then applied to ELISA plates, preincubated for 15 min with 50 nM of biotinylated target proteins at RT. Plates with added phages were incubated for 15 min at RT and washed 3-times with 1×PBST. The washing step was followed by 30 min incubation with HRP-conjugated anti-M13 mouse monoclonal antibody diluted in PBST in 1:5000 ratio. Excess antibody was washed away with 1×PBST and plates were developed using TMB substrate, quenched with 1.0 M HCl and the signal was determined by absorbance measurement (A_450_).

Protein based single-point ELISA was performed to confirm binding of generated unique sABs to their cognate antigens. Immobilized on ELISA plate target (50 nM) was incubated with 200 nM of the purified sABs for 15 min. The plates were then washed and incubated with a secondary HRP-conjugated anti-human F(ab’)_2_ monoclonal antibody (1:5000 dilution in PBST). The plates were washed, developed with TMB substrate and quenched using 1.0 M HCl, and absorbance (A_450_) signal was measured.

### Cloning, Overexpression and Purification of sABs

Phage ELISA results were used to select sAB clones that were sequenced at DNA Sequencing Facility at The University of Chicago. In-fusion cloning (66) was used to reformat unique sABs clones into pRH2.2, an IPTG inducible vector for bacterial expression.

*E. coli* BL21 (Gold) cells were transformed with sequence-verified sAB plasmids. Cultures were grown in 2YT media supplemented with 100 μg/mL at 37°C until they reach OD_600_ = 0.8, when they were induced with 1 mM IPTG. The culture was continued for 4.5h at 37°C and cells were harvested by centrifugation. Cell pellets were resuspended in 20 mM HEPES pH = 7.5, 200 mM NaCl, 1 mM PMSF, 1 μg ml^-1^ DNase I, and lysed by ultrasonication. The cell lysate was incubated at 60°C for 30 min. Heat-treated lysate was centrifuged at 50,000 *x g* to remove cellular debris, filtered through a 0.22 μm filter and loaded onto a HiTrap protein L (GE Healtchcare) column pre-equilibrated with 20 mM HEPES pH 7.5 and 500 mM NaCl. The column was washed with 20 mM HEPES pH 7.5 and 500 mM NaCl and sABs were eluted with 0.1 M acetic acid. Protein containing fractions were loaded directly onto an ion-exchange Resource S column pre-equilibrated with 50 mM NaOAc pH 5.0 and washed with the equilibration buffer. sABs elution was performed with a linear gradient 0–50% of 50 mM NaOAc pH 5.0 with 2 M NaCl. Purified sABs were dialyzed overnight against 20 mM HEPES Ph 7.5 with 150 mM NaCl. The quality of purified sABs was analyzed by SDS-PAGE.

### Binding Kinetics by SPR

SPR experiments were performed at RT using BIACORE 3000 (GE Healthcare) instrument. Targets were immobilized onto a nitrilotriacetic acid (NTA) sensor chip via His-tag. For kinetic experiments, 2-fold serial dilutions of the sAB were injected following ligand immobilization on the sensor chip. For kinetic assay six dilutions of the sAB were tested.

### Flow cytometry

HEK293T cells were seeded in 6-well plates with DMEM supplemented with 10% (v/v) FBS. After 24 h, cells reached 50-60% confluence and were transfected with 1 µg of ADGRL3 or ADGRL1 (41) or EV using 6 µl of transfection reagent LipoD293T. After 48 h, cells were detached with citric saline solution and washed with 0.1 % BSA in PBS. The pellets were incubated with synthetic antibodies 30 mins at 4 C, washed twice with 0.1 % BSA in PBS and incubated with fluorescent-tagged secondary antibody (Alexa 647) for 30 min at 4 C and washed three times and fixed in 1% paraformaldehyde. Flow cytometry data were collected on BD Accuri C6 flow cytometer and analyzed in FlowJo.

### Serum Response Element Luciferase Assays

HEK293T cells were seeded on a 96-well flat-bottom plate precoated with 0.5% gelatin and grown until 50-60% confluent in DMEM supplemented with 10% (v/v) FBS. Cells were then co-transfected with ADGRL3/ADGRL1 constructs (1 ng well^-1^ for ADGRL3; ng well^-1^ for ADGRL1) (41), Dual-Glo luciferase reporter plasmid (20 ng well^-1^) (21), using 0.3 μL LipoD293T (SL100668; SignaGen Laboratories). DNA levels were balanced among transfections by addition of the empty pCMV5 vector to 100 ng total DNA. 18 h after transfection media was aspirated and replaced with DMEM without FBS. When sABs were tested, 2 µM of sAB was added 5h after start of the serum-starvation. After 10h of serum starvation, the media was removed and cells were lysed using Dual-Glo Luciferase Assay System from Promega and firefly and renilla luciferase signals were measured using a Synergy HTX (BioTek) luminescence plate reader. The firefly:renilla ratio for each well was calculated and normalized to empty vector.

### cAMP Assay

HEK293 cells were seeded in 6-well plates with DMEM supplemented with 10% (v/v) FBS. After 24 h, cells reached 50-60% confluence and were transfected with 350 ng of ADGRL3 + 350 ng 22F Glosensor reporter plasmid (E2301; Promega – a gift from R. Lefkowitz lab) + 9 ng of β2-adrenergic receptor + 2.8 uL of transfection reagent Fugene 6. After 24 h, cells were detached and seeded at 5× 10 ^4^ per well in white flat bottom 96-well plate. After another 24 h, the media was replaced with 100 ul Opti-MEM I Reduced-Serum Medium (31985070, Life Technologies) and incubated for 30 min. Then, 1 uL Glosensor substrate and 11 uL FBS were added to each well. Basal cAMP signal was measured after 20 min of equilibration time. Next cells were treated with 2 uM of sAB for 5 min and then activated with 50 nM isoproterenol. Measurements were done using Synergy HTX BioTek plate reader at 25 C.

### Formation of ADGRL3_Lec_/LK30 Complex

ADGRL3/sAB complex was formed by mixing 1.5-fold molar excess of the Lec domain with the LK30 sAB and 30 min incubation on ice. Next, the complex was subjected to size-exclusion chromatography on a Superdex 200 10/300 column pre-equilibrated with 30 mM HEPES pH 7.5 with 150 mM NaCl. Formation of the ADGRL3_Lec_/LK30 complex was determined by retention volume analysis of the complex with respect to that of target alone and co-elution of the individual components on SDS-PAGE.

### X-ray crystallography

Purified ADGRL3_Lec_/LK30 complex was crystallized using hanging-drop vapor diffusion at 5 mg mL−1 in 100 mM sodium acetate (pH=4.5), 150 mM ammonium sulfate and 20% (w/v) PEG 4000. Crystals were frozen in mother liquor with the addition of 20% glycerol. Data were collected to 2.65 Å at the Advanced Photon Source at Argonne National Laboratory (beamline 23-ID-B). The datasets were auto processed using the GM/CA beamline GMCAproc protocol, which employs XDS (67) and pointless (68) to index, integrate, scale, and merge the data. Initial phases were determined by molecular replacement using PHASER (69) in the CCP4 suite (70), utilizing the available human Lec domain structure (PDB ID: 6VHH) (39) and sAB structure (PDB ID: 4XWO) (71) as search models. The structure of the complex was obtained in space group P2_1_2_1_2_1_ with two ADGRL3_Lec_/LK30 complexes in the asymmetric unit. Initial rounds of refinement and model building were performed with REFMAC5 using NCS restraints. Next, phenix.refine (72) was used without NCS restraints but with reference model restraints, using high resolution structures of the Lec domain (PDB 5AFB) (73) and sAB (PDB 5UCB). Final rounds of model building and refinement were performed in phenix.refine without reference model restraints or NCS restraints. Final refinement parameters were rigid body refinement with individual B-factors, TLS refinement, and optimization of stereochemistry and ADP weighting.

### LK30 binding to ADGRL/TEN and ADGRL/FLRT complexes

For ADGRL3/TEN2/LK30 complex test, ECRs of ADGRL3 and TEN2 were co-expressed in High Five insect cells, purified by Ni-NTA affinity chromatography and subjected to SEC. Purified fractions were pulled and 2-fold molar excess of LK30 was added, followed by 30 min incubation on ice. Next, the mixture was subjected to SEC on a Superose 6 10/300 column pre-equilibrated with 30 mM HEPES pH 7.5 with 150 mM NaCl.

For AGRL3/FLRT3/LK30 tests, ECRs of ADGRL3 and FLRT3 were expressed in High Five insect cells. After purification of individual proteins, the complex was formed by mixing proteins in 1:1 molar ratio at room temperature for 30 min, followed by SEC purification. Then, 2-fold molar excess of LK30 was added, the mixture was incubated for 30 min on ice and injected on Superdex 200 10/300 column pre-equilibrated with 30 mM HEPES pH 7.5 with 150 mM NaCl.

### Cell Aggregation Assay

HEK293T cells were seeded in 6-well plate containing 2.5 mL of DMEM media supplemented with 10% FBS and incubated overnight at 37 °C. When cells reached 80% confluency, they were then co-transfected with 2 µg of either pCMV5 + GFP, pCMV ADGRL3 + GFP, TEN2 + dsRed, or FLRT3 + dsRed using 4 μL LipoD293T (SL100668; SignaGen Laboratories). Two days after transfection, the media was aspirated, cells were washed with 1xPBS and detached with 1xPBS containing 1 mM EGTA and supplemented with 15 µL of 1 mg/20 µL DNAse (Sigma, D5025). Cells were resuspended by pipetting to a create single-cell suspensions, transferred to an Eppendorf tube and additional 15 µL of DNAse solution was added to each sample. Seventy µL of cells expressing indicated constructs were mixed in 1:1 ratio in a one well of a non-coated 24-well plate containing 340 µL of Incubation Solution (DMEM supplemented with 10% FBS, 10 mM CaCl_2_ and 10 mM MgCl_2_). Forty µL of either sAB LK30 (to final concentration of 5 µM) or 1xHBS was added to the mixture, and cells were then placed on a shaker at 120 rpm at 27 °C for 10 min and imaged using a Leica Fluorescent Microscope with a 5x objective. Aggregation index at time=10 was calculated using ImageJ. A value for particle area of 1-2 cells was set as a threshold based on negative control values. The aggregation index was calculated by dividing the area of particles exceeding this threshold by the total area occupied by all particles in the individual fields.

## ACKNOWLEDGEMENTS

We thank Engin Özkan for the use of flow cytometer and all members of the Araç lab for helpful discussions. We also thank the staff at the Advanced Photon Source (APS) at Argonne National Labs (ANL). GM/CA@APS has been funded by the National Cancer Institute (ACB-12002) and the National Institute of General Medical Sciences (AGM-12006, P30GM138396). This research used resources of the Advanced Photon Source, a U.S. Department of Energy (DOE) Office of Science User Facility operated for the DOE Office of Science by Argonne National Laboratory under Contract No. DE-AC02-06CH11357. The Eiger 16M detector at GM/CA-XSD was funded by NIH grant S10 OD012289. This work was supported by grants R01 GM120322 (to D.A.), R01 GM134035-01 (to D.A.) and GM117372 (to A.A.K.).

## AUTHOR CONTRIBUTIONS

S.P.K. and D.A. designed all experiments and interpreted results. S.P.K. expressed and purified all proteins (with assistance from B.A. and K.L.). P.D. carried out phage display selection and sABs characterization. S.P.K. and J.M.A. performed flow cytometry experiments and cell-based signaling assays. and S.P.K. performed crystallography experiments (with assistance from B.A.) and structure determination. J.L. participated in cell-cell aggregation experiments. S.P.K. and D.A. wrote the manuscript with assistance from A.A.K., S.J.B. and P.D.. D.A. and A.A.K. supervised the project.

---

**Supplementary Figure 1.**
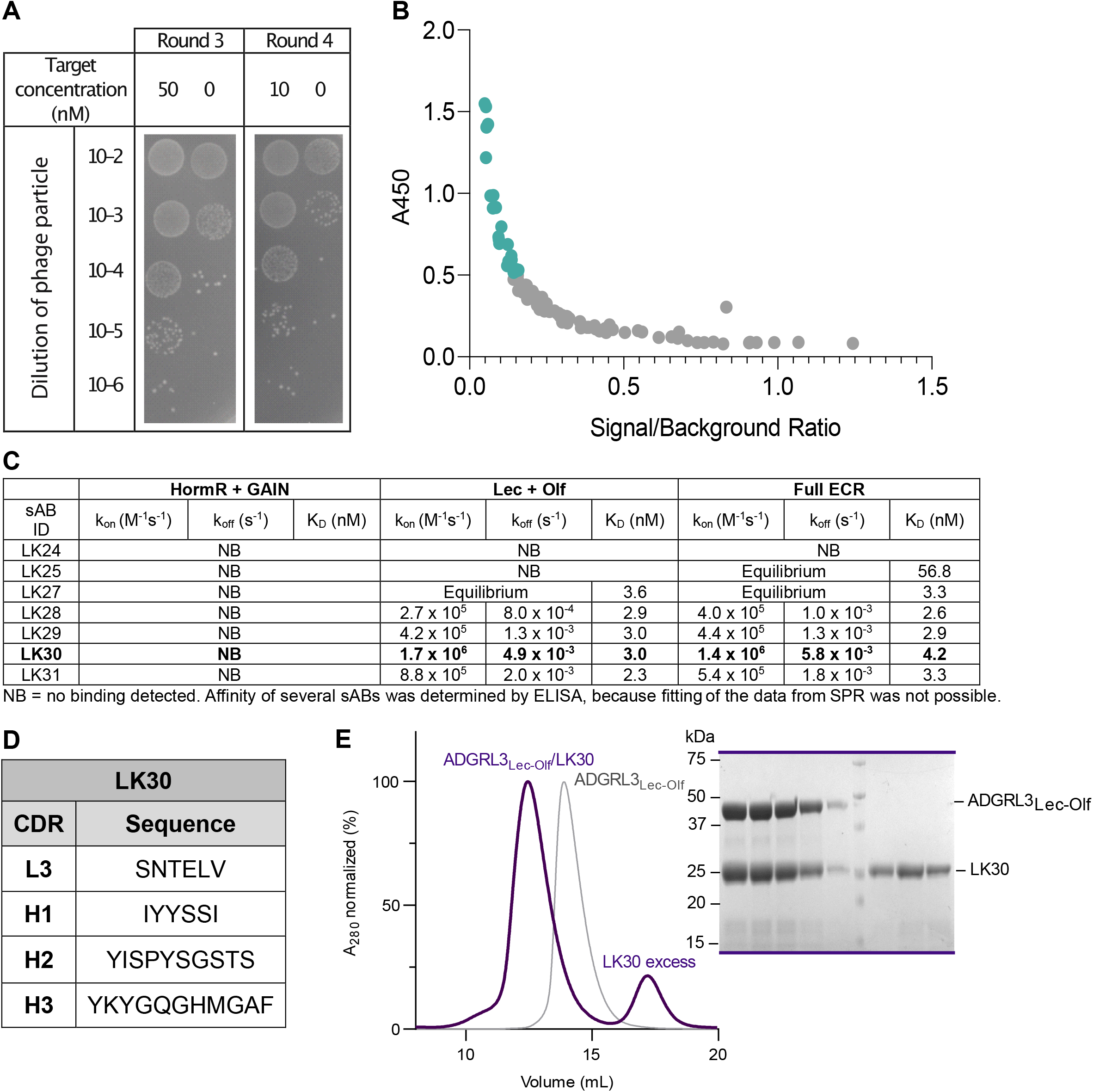
Development and characterization of sABs against ADGRL3. **A**.Enrichment of the target-specific binders after third and fourth round of antibody selection for full ECR of ADGRL3. **B**.Binders obtained after four rounds of phage display antibody selection were screened by a single-point phage ELISA. Clones marked in cyan were further characterized. **C**. Binding kinetics of unique sABs developed against ECR of ADGRL3. SPR values of LK30 binding to either purified full-length ECR of ADGRL3, Lec/Olf fragment, and HormR/GAIN fragment. **D**. Sequences of LK30 CDRs. **E**. SEC profiles and SDS-PAGE analyses of the LK30 complex with Lec-Olf fragment of ADGRL3.

**Supplementary Figure 2.**
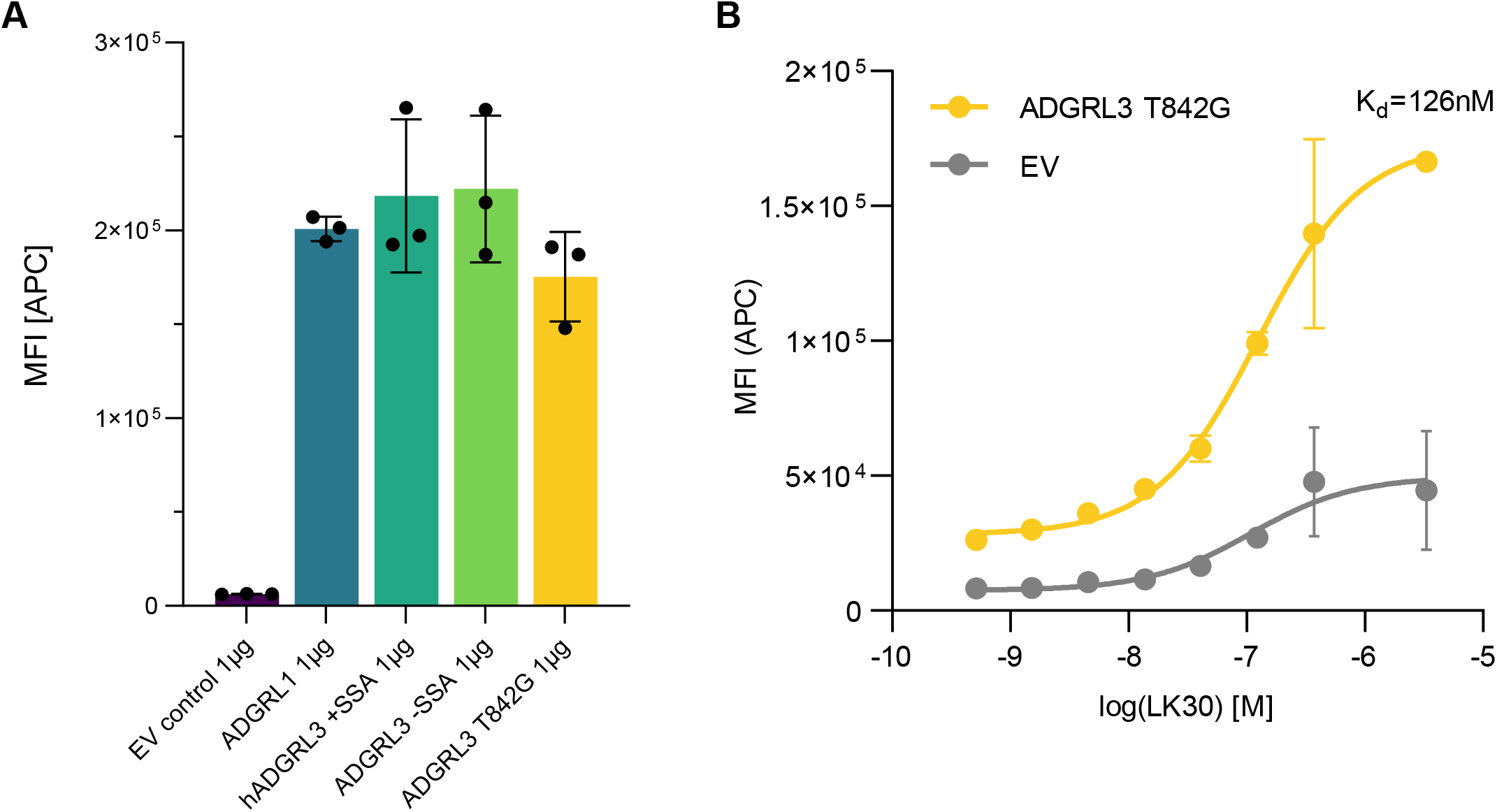
sAB LK30 binds to the ADGRL3 autoproteolysis mutant. **A**. Cell surface expression of all constructs was tested by transfecting HEK293T cells with equal amount of construct DNA, and measuring the protein expression using anti-FLAG antibody by flow cytometry. Error bars indicated SD (n = 3). **B**. Binding of LK30 to the ADGRL3 T842G autoproteolysis mutant was measured on HEK293T cells by flow cytometry. K_D_ value of LK30 binding was determined to be 126 nM. Error bars indicated SD (n = 3).

**Supplementary Figure 3.**
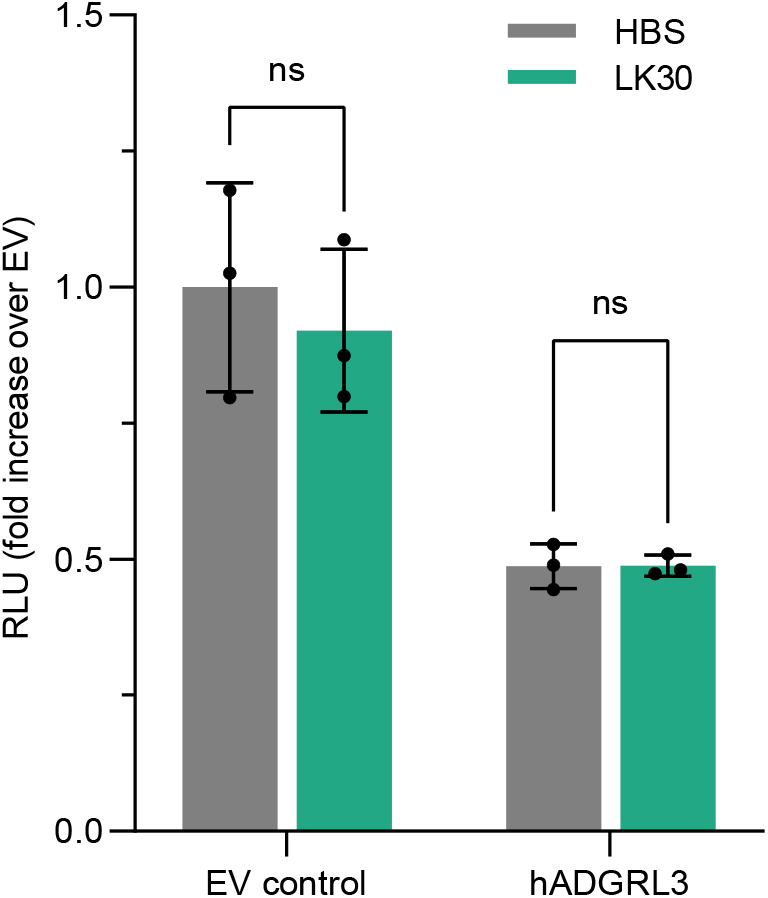
LK30 does not affect ADGRL3 cAMP signaling. ADGRL3 signaling measured by the cAMP assay in the presence of 2 µM of LK30. Data are shown as a fold decrease over EV. RLU, relative luminescence units. Error bars indicated SD (n = 3).

**Supplementary Figure 4.**
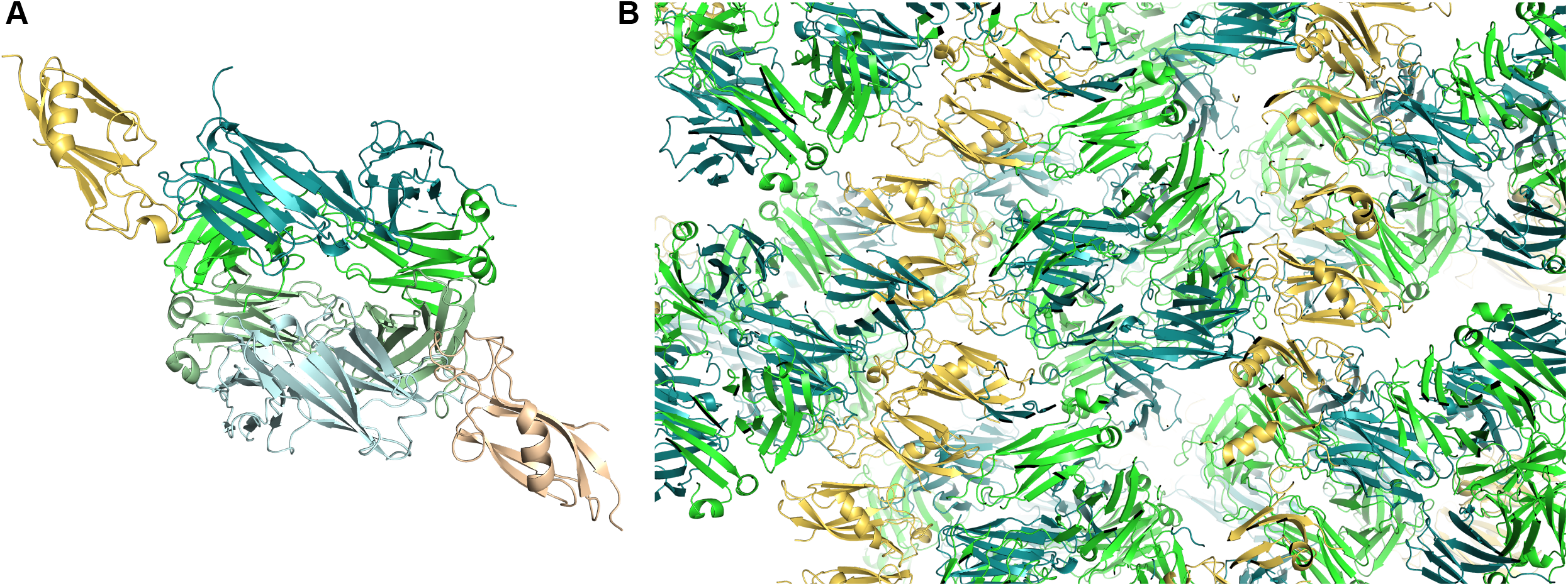
Crystal structure of ADGRL3/LK30 complex. **A**. The structure of the complex was obtained in space group P2_1_2_1_2_1_ with two ADGR3_Lec_/LK30 complexes in the asymmetric unit. **B**. Analysis of the crystal packing shows the contacts in the crystal lattice are mediated primarily by the heavy chain (HC) and light chain (LC) of LK30 and Lec domain of ADGRL3. ADGRL3Lec is colored yellow while the HC and LC of LK30 are colored green.

**Supplementary Figure 5.**
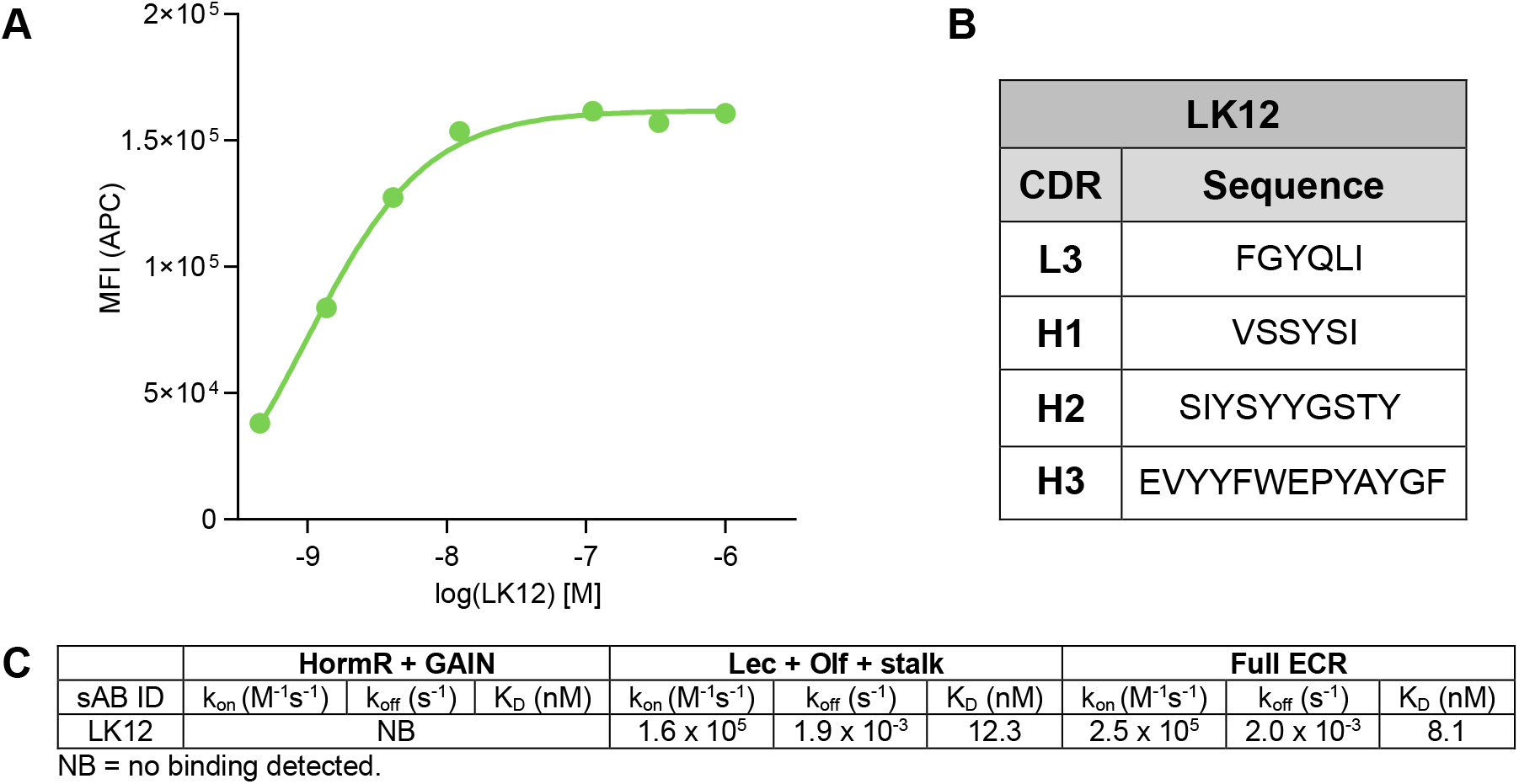
Characterization of ADGRL3 binder sAB LK12. **A**. Binding activity of LK12 to the receptor was measured using HEK293T cells expressing full-length ADGRL3 by flow cytometry. **B**. Sequences of LK12 CDRs. **C**. Binding kinetics of unique sABs developed against ECR of ADGRL3. SPR values of LK30 binding to either purified full-length ECR of ADGRL3, Lec/Olf fragment, and HormR/GAIN fragment.

